# Towards a database capturing chromosome structure and function: symbols and syntax

**DOI:** 10.64898/2026.05.14.724942

**Authors:** Peter R. Cook, Davide Marenduzzo, Zahra Valei

**Affiliations:** The Sir William Dunn School of Pathology, University of Oxford, South Parks Road, Oxford OX1 3RE, UK; SUPA, School of Physics and Astronomy, University of Edinburgh, Peter Guthrie Tait Road, Edinburgh EH9 3FD, UK

**Keywords:** chromosome organization, transcription, database

## Abstract

Existing databases of interphase chromosome conformations typically store three-dimensional coordinates of genomic segments. However, since interphase chromatin is highly dynamic, such databases are dominated by transient configurations and unstructured regions, whose positions vary continuously between cells and over time, unlike folded proteins such as globin, which adopt similar structures in every cell. These drawbacks motivated the inception of a database based on ‘strion’ (a portmanteau of a string capturing *str*ucture and funct*ion*). A strion concisely describes the structure and activity of all transcription units in one cell, by retaining only functionally relevant positional information. Sets of strions describing structures in different cells sampled at different times are compiled into a ‘super-strion’. Then, 46 super-strions summarise the range of structure and activity of a human cell type, including information on all transcription units, how often each co-fires and co-clusters with others in transcription factories/hubs, enhancer interactomes and small-world expression networks.

**Graphical abstract:** 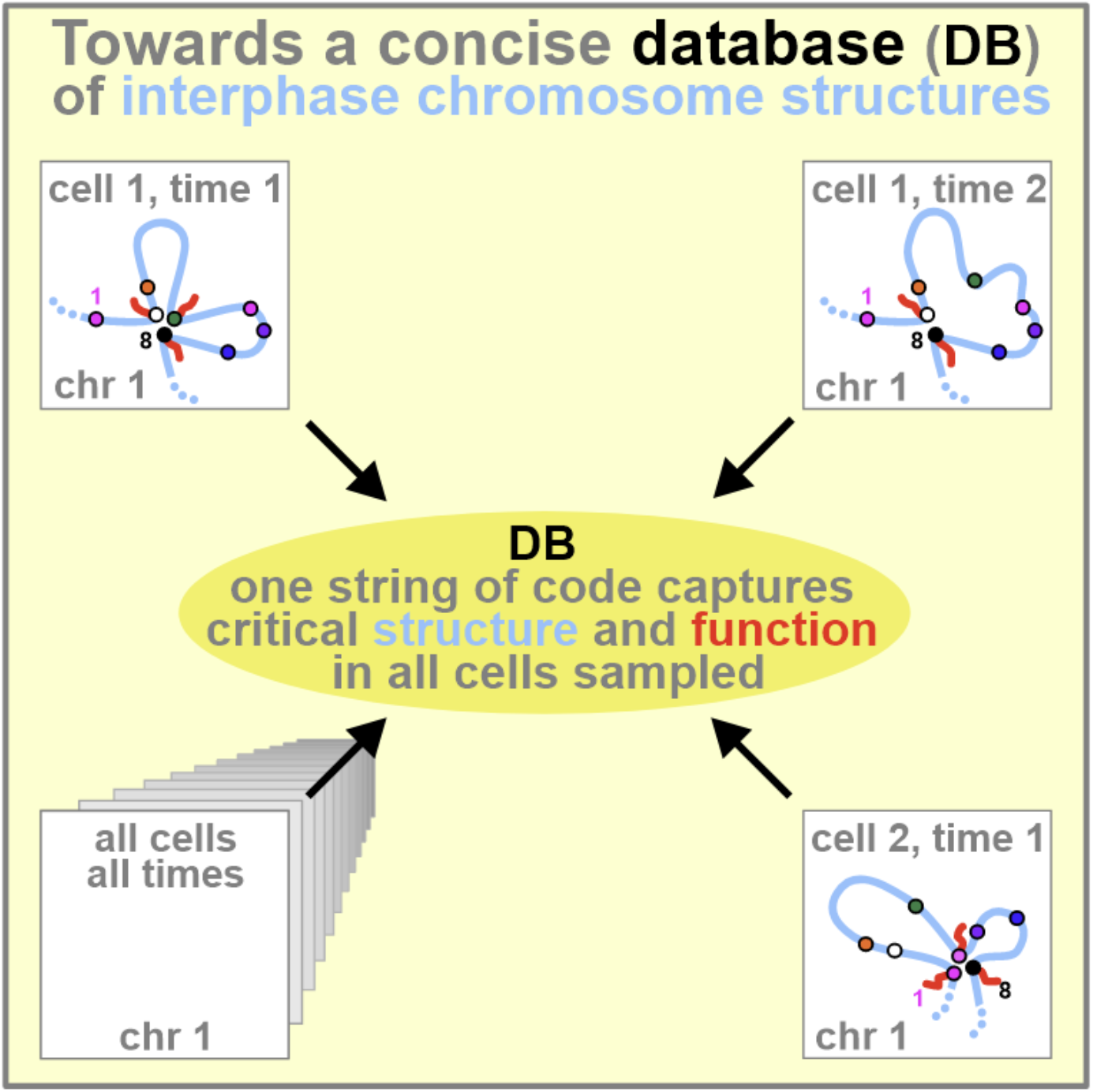

## Introduction

Structures of interphase chromosomes are being deposited in increasing numbers in databases (Dekker *et al*., 2026). This raises a natural question: what kind of information should such databases provide? By analogy to the Protein Data Bank (PDB; Berman *et al*., 2000)—which lists the coordinates of atoms in proteins—the answer seems obvious. Chromosome databases should list the three-dimensional (3D) positions of individual bases, and ultimately their constituent atoms. Whilst current chromosome databases list 3D coordinates of large genomic segments (e.g., Oluwadare *et al*., 2019; Oluwadare *et al*., 2020; Contessoto *et al*., 2021; Chiang *et al*., 2022, Chiang *et al*., 2024), the clear intention is to provide coordinates at the level of individual bases and atoms.

We suggest a PDB-like database of chromosome structures has major drawbacks. First, each atom in the PDB has a fixed position relative to others in the protein, but in most living cells, the majority of chromosome segments change from moment to moment (e.g., Lerner *et al*., 2020). This is because most base pairs reside in loops, with only the anchor points remaining (transiently) static (Misteli, 2020). For example, loops in HeLa cells show a broad distribution of contour lengths, with a mean of ∼86 kbp (Jackson *et al*., 1990). Assuming that anchored segments are ∼20 bp, <<0.1% of the genome would then be static. In other words, nearly all information in current chromosome databases concerns unstructured regions, which are normally excluded from the PDB. Second, it is unlikely that the structure of any human chromosome deposited now or in the future in a PDB-like database will ever be found in the same cell later. This is because the structure is in continual flux and the number of possible conformations is extremely large. Third, chromosome databases must soon become enormous, due to the relative sizes of chromosomes and proteins. There are 60-68 atoms in a base pair, compared to 20-30 in an amino acid. As a result, a 100 Mbp chromosome has at least a thousand-fold more atoms than a 50 kD protein (∼6.4×10^9^ compared to ∼10^4^). Moreover, in contrast with the single string of code defining a protein in all cell types of a species, many strings will be needed just to begin sampling the structural range adopted by one chromosome in a single cell over a few minutes. Even more strings will be required for the same chromosome in other cells of the same type, and still more for cells of different types (Hatton *et al*., 2023). Fourth, structural data are derived from diverse experimental approaches and are typically stored in incompatible formats. Finally, both the PDB and current chromosome databases largely lack direct functional annotation.

Despite these challenges, existing data consistently point to a key feature: a minority of genomic segments, which are loop anchors, are those most closely associated with gene regulation, as they bring specific genomic elements into spatial proximity. This observation and the combination of drawbacks mentioned above motivated the inception of the database proposed here.

Our purpose is to open discussion of what one might like to see in a chromosome database. As a first step, we propose symbols and syntax suitable for code that concisely captures the ever-changing interplay of conformation and function. Here, function refers to transcriptional activity, which is currently best determined using GRO-/PRO-seq (global/precision run-on sequencing) or its variants (Kwak *et al*., 2013; Jordán-Pla *et al*., 2019). This code is deliberately designed to describe the 3D coordinates of just the (transiently) structured minority, whilst retaining sufficient 3D information to mark the positions of every active (or potentially active) transcription unit (TU) in the cell type studied. It also tracks the general path of each chromosome through space (and thus the rough limits of chromosome territories) and the locations of loop anchors. It additionally provides functional information on how often each TU fires, plus associated enhancer interactomes and small-world transcriptional networks. In this way, this is a stripped-down structural database that incorporates functional data. We suggest that it should prove useful both now, with incomplete and sparse data, and in the future, when atomic resolution is achieved. It can be designed to complement existing larger, technique-specific databases through use of the Common Coordinate Framework (HuBMAP Consortium, 2019).

### Outline

We begin by describing a “database of the future” (which we will call the complete database) that would be useful when the 3D coordinates of every base pair in every chromosome in each cell of a given cell type are known. We then discuss variants suitable for use with the sparse data currently available. For illustrative purposes, we mainly focus on human and mouse chromosomes and TUs transcribed by RNA polymerase II, along with chromosomal contacts detected by Hi-C (a high-throughput variant of 3C, chromosome conformation capture; Lieberman-Aiden *et al*., 2009) and simulations performed with coarse-grained molecular dynamics (Di Pierro *et al*., 2018; Chiang *et al*., 2024; Yildirim *et al*., 2022; Giorgetti *et al*., 2014; Tiana and Giorgetti, 2018; Bianco et al., 2018). This focus reflects the abundance of available data.

However, our approach is general and can be applied to data from any organism, obtained using other methods like pore-C (an analogue of Hi-C using nanopore technology; Deshpande *et al*., 2022; Dotson *et al*., 2022), GAM (genome architecture mapping; Beagrie *et al*., 2017; Winick-Ng *et al*., 2021), SPRITE (split-pool recognition of interactions by tag extension; Quinodoz *et al*., 2018), RCMC (region-capture Micro-C; Goel *et al*., 2023), multiplexed FISH (fluorescence *in situ* hybridization; Bintu *et al*., 2018; Nir *et al*. 2018; Shah *et al*, 2018), and ChromExM (chromatin expansion microscopy; Pownall *et al*., 2023). The framework can also be extended to include TUs transcribed by other RNA polymerases, and additional motifs or landmarks such as histone marks, CCTF-binding sites, bound cohesins, centromeres and telomeres.

### Active transcription units as motifs central to chromosome structure and function

We use the term ‘TU’ to refer to any genic or non-genic transcription unit that is active or potentially active in the cell type considered. This definition includes enhancers, which are now widely regarded as non-genic promoters that function through transcription in close spatial proximity to their target genes (Andersson *et al*., 2014; Furlong and Levine, 2018; Schoenfelder and Fraser, 2019; Andersson and Sandelin, 2020; Goel *et al*., 2023; Negro *et al*., 2024). In humans, enhancers outnumber genes by approximately 10:1 (Andersson *et al*., 2014). For example, when chromosome HSA14 in HUVECs (length ∼10^7^ Mbp) is coarse-grained into 1 kbp segments, there are ∼3000 segments that encode active or potentially active TUs but only ∼350 are genic ones.

Active TUs will be the central motifs in our proposed database for three main reasons. First, they play key roles in determining both chromosome structure and function (Misteli, 2020; Rippe and Papantonis, 2025; Dekker *et al*., 2026), and provide a natural link between spatial organisation and transcriptional output (Brackley *et al*., 2021; Negro *et al*., 2024). Second, high-resolution measurements of chromosomal contacts—obtained using RCMC—show that interactions involving active TUs far outnumber others; for example, transcribed promoters and enhancers stabilize 67– 74% contacts, compared with only ∼4% involving CTCF and cohesin (Goel *et al*., 2023). Similar conclusions are supported by bacterial Hi-C, where active polymerases anchor most loops (Bignaud *et al*., 2024).

Third, it is increasingly accepted that TUs fire after attaching to clusters of active polymerases— referred to variously as hubs, condensates, (micro)compartments, or transcription factories (Goel *et al*., 2023; Harris *et al*, 2023; Pownall *et al*, 2023; Negro *et al*., 2024; Rippe and Papantonis, 2025). Early studies also showed that at least 92% of all nascent polymerase II transcripts were made in such clusters (reviewed by Negro *et al*., 2024). Therefore, such clusters contain two or more active TUs, each providing an anchor point to yield some contacts detected by Hi-C. In our cartoons, anchors are indicated by a red circle with an attached nascent RNA to indicate present or future activity. However, many anchors may instead involve transcription factors transiently bound to potentially-active promoters rather than active polymerases (Cook & Marenduzzo, 2018; Goel *et al*., 2023; Negro *et al*., 2024).

### Hi-C and structure determination

Hi-C is the most widely used method for determining chromosome structure. It provides information on intra- and extra-chromosomal contacts (i.e., *cis* and *trans* contacts), plus 2-way and higher-order ones (Dekker *et al*., 2026). However, many Hi-C pipelines discard higher-order and *trans* contacts. A single Hi-C experiment also fails to detect most human loops shorter than ∼200,000 bp (Rowley *et al*., 2020), which contrasts with median values of <100 kbp found using earlier methods (e.g., Cook and Brazell, 1975; Jackson *et al*., 1990; Nickerson, 2001). Additionally, Hi-C underestimates the presence of the transcription machinery at anchors (i.e., active polymerases and transcription factors) and the number of clustered contacts (O’Sullivan *et al*., 2013; Quinodoz *et al*., 2018; Beagrie *et al*., 2023; Zhang *et al*., 2023). Finally, it does not initially provide 3D coordinates.

In constructing the complete database, we therefore apply a simple principle: we retain only contacts involving TUs, and prioritise higher-order interactions where multiple TUs co-localise. This reflects the view that such contacts are the most informative about chromosome structure and function.

This approach can be illustrated using two chromosomal segments that yield Hi-C contacts (**Fig. 1A**). Most DNA in these segments is mobile and unstructured, with only a small number of TUs (e.g., α, 1, and 5) acting as transient anchors. These anchors define a cluster, which we assume determines both local structure and transcriptional activity (reviewed by Negro *et al*., 2024). The majority of Hi-C contacts (e.g., b–c) arise from chance encounters between unstructured regions within loops and are therefore discarded (**Fig. 1A**; black line). By contrast, contacts involving TUs, such as pairwise (2–d, 3–4) or higher-order (α–1–5) interactions, are retained (**Fig. 1A**; purple lines).

**Figure 1.**
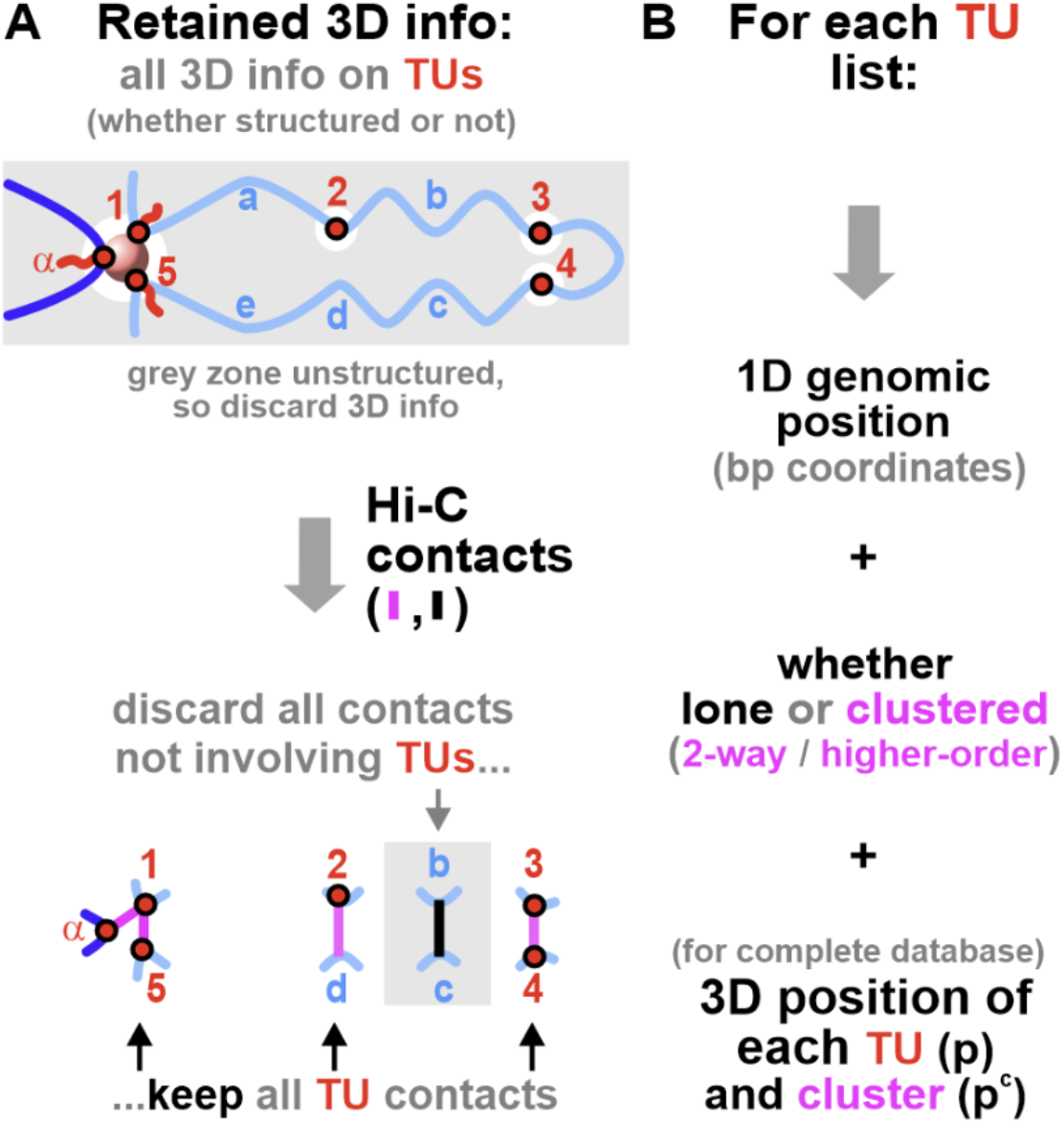
Data required for our databases illustrated using those from Hi-C. **A**. Active TUs 1 and 5 (on one chromosome), and *α* (on another) form what we call a ‘cluster’ that anchors 2 different chromosomes (light and dark blue). Parts of chromosomes far from anchors are unstructured. Contacts: we keep only those involving one or more TUs (purple lines). All other contacts, such as those between non-TU DNA (black line), are discarded. 3D information: we retain positions of all TUs—whether in structured regions (like 1,5, *α*, anchored to the pink hub) or in unstructured regions (like 2, 3, 4). **B**. Some required information.

Polymer models, such as the fractal globule, enable prediction that most pairwise contacts arise from random encounters and that higher-order contacts are comparatively rare. Consequently, repeated detection of a contact like *α*-1-5 (or a higher-order one) across cells is unlikely to occur by chance (**Table I**). Higher-order contacts therefore contain the most information on the original structure and its function. They appear consistently in raw data from Hi-C (Olivares-Chauvet P *et al*., 2016), pore-C (Dotson *et al*., 2022), GAM (Beagrie *et al*., 2023), SPRITE (Quinodoz *et al*., 2018), and simulations (e.g., Brackley *et al*., 2021; Bianco *et al*., 2018), but have not been studied as extensively as 2-way ones. Nevertheless, the highest-resolution RCMC and Hi-C data still reveal the prior existence of higher-order forms, despite selecting only 2-way *cis* contacts for analysis. This is indicated by (micro)compartments making numerous off-diagonal and focal/nested contacts between TUs (Goel *et al*., 2023; Harris *et al*., 2023; Li *et al*., 2023).

**Table I.** Probabilities of chance contacts between chromosomal segments, and the number of TU multi-way contacts recorded in a population of 10^6^ cells which is needed for statistical significance. Probability of contacts are computed assuming: (i) segments are 1 kbp in length, (ii) nucleoplasmic diameter is 10 *µm*, (iii) the threshold for contacts between two segments is 20 nm centre-to-centre distance, (iv) genome size is 6 Gbp, and (v) a typical 100 Mbp chromosome occupies ∼1/50 of nucleoplasmic volume. For the fractal globule case, we assume the probability of contact between two segments 100 kbp apart is 1/s=0.01. Number of contacts needed for significance are computed by assuming there are 200000 TUs in the diploid genome, and 3000 TUs in a 100 Mbp chromosome. Significance requires that the “false discovery rate”, or the number of 2-way or 3-way TU contacts expected randomly is at least <0.01. These estimates show that multi-way TUs observed multiple times are highly likely to be statistically significant (and functionally relevant).

| Type of contact | Intra-chromosomal |  | Inter-chromosomal |
| --- | --- | --- | --- |
|  | Chromosomal segments randomly assorted | Assuming chromosome is fractal globule, and contact 100 kb away | All segments in all chromosomes randomly assorted |
|  | Probability that contact between segments occurs by chance |  |  |
| Specific segments: two-way | $5 \times 10^{-6}$ | $10^{-2}$ | $10^{-7}$ |
| Specific segments: three-way | $2.5 \times 10^{-11}$ | $10^{-4}$ | $10^{-14}$ |
| | Number of contacts seen in $10^6$ cells needed for significance | | |
| TUs two-way | 24 | ~10400 | 8 |
| TUs three-way | 3 | ~130 | 3 |

Retaining information on TUs located within otherwise unstructured regions (e.g., 2, 3, and 4 in Fig. 1A) allows the general path of each chromosome and the degree of intermingling between chromosomes to be tracked. However, we will see that excluding non-TU contacts has little impact on the primary functional uses of the database.

Finally, although raw Hi-C data yields more *cis* contacts (like 1-5 in **Fig. 1A**) than *trans* ones (like 1-*α* in **Fig. 1A**), especially when applied with in-nucleus ligation (Nagano *et al*., 2015), other methods uncover the opposite pattern. For example, after pulling down RNA polymerase II, ChIA-PET recovers more *trans* than *cis* contacts (Li *et al*., 2012; Papantonis *et al*., 2012). Similarly, ligation-free single-cell SPRITE yields 54% *trans* contacts, compared to 6% with single-cell Hi-C (Arrastia *et al*., 2022). Additionally, intron-seqFISH shows that 82.4% nascent genic RNAs in mouse ES cells lie <500 nm from a *trans* counterpart (Shah *et al*, 2018). These observations indicate that *trans* contacts between active TUs are common and should be incorporated in the database.

### Information required for TU-based chromosome databases

In all our databases, chromosome structure is represented in terms of TUs and their spatial organisation. For each TU, we record two key pieces of information: genomic position (in linear base-pair coordinates) and whether the TU is alone or clustered (**Fig. 1B**).

A cluster is defined as one or more TU that lie within a specified distance of each other or another DNA segment. This distance depends on the detection method. In Hi-C, clusters are detected through 2-way and higher-order contacts (with an effective interaction range sufficiently short to permit ligation), whereas in simulations they are defined by an explicit distance threshold between polymer beads. Empirical estimates can be based on well-characterised transcriptional clusters (factories) in HeLa cells, which have diameters of ∼50–175 nm (mean ∼90 nm) (Cook and Marenduzzo, 2018; Negro *et al*., 2024). More precise, technique-specific values can be incorporated as they become available. Each cluster is associated with a contact list (c_list) specifying its associated TUs.

In a complete database, we additionally record the 3D position of each TU (abbreviated as p with a superscript to be described) and each cluster (abbreviated as p^c^, determined by triangulation from positions of constituent TUs; **Fig. 1B**). Obtaining such 3D coordinates remains technically challenging, although they are available from simulations (e.g., Brackley *et al*., 2021; Harris *et al*., 2023). Where possible, the positions of confining nuclear structures (e.g., lamina, nucleoli) can also be included, approximated initially by simple geometries such as spheres or ellipsoids. However, these data are currently sparse and are not considered further here.

### TUs are members of different small-world networks

A typical TU is inactive for most of the time as it diffuses through the nucleoplasm (Larsson *et al*., 2019). It only joins the structured minority when it binds to a cluster, and so is likely to be transcribed (Negro *et al*., 2024). In mammalian cells, polymerase II clusters typically anchor ∼10 loops to yield inter-TU contacts detectable by Hi-C (Goel *et al*., 2023; Negro *et al*., 2024; Rippe and Papantonis, 2025). TUs that bind the same transcription factors often co-localise in the same clusters. As a result, different clusters specialise in transcribing different groups of TUs that are parts of different small-world networks (Cook & Marenduzzo, 2018; Chiang *et al*., 2022; Dotson *et al*., 2022; Negro *et al*., 2024), consistent with the organisation of gene-expression networks (e.g., van Noort *et al*., 2004; Badia-i-Mompel *et al*., 2023). Different RNA polymerases, I, II, and III, are similarly associated with distinct clusters (e.g., Pombo *et al*., 1999), and genes binding different factors co-cluster when transcribed by polymerase II (e.g., ERα, KLF1, NFκB, and TFEC; Fullwood *et al*., 2009; Schoenfelder *et al*., 2010; Papantonis *et al*., 2012; Dotson *et al*., 2022). For simplicity, our schematics illustrate only a small number of cluster types (Fig. 2), although a much richer diversity is likely to exist (Semeraro *et al*., 2025).

**Figure 2.**
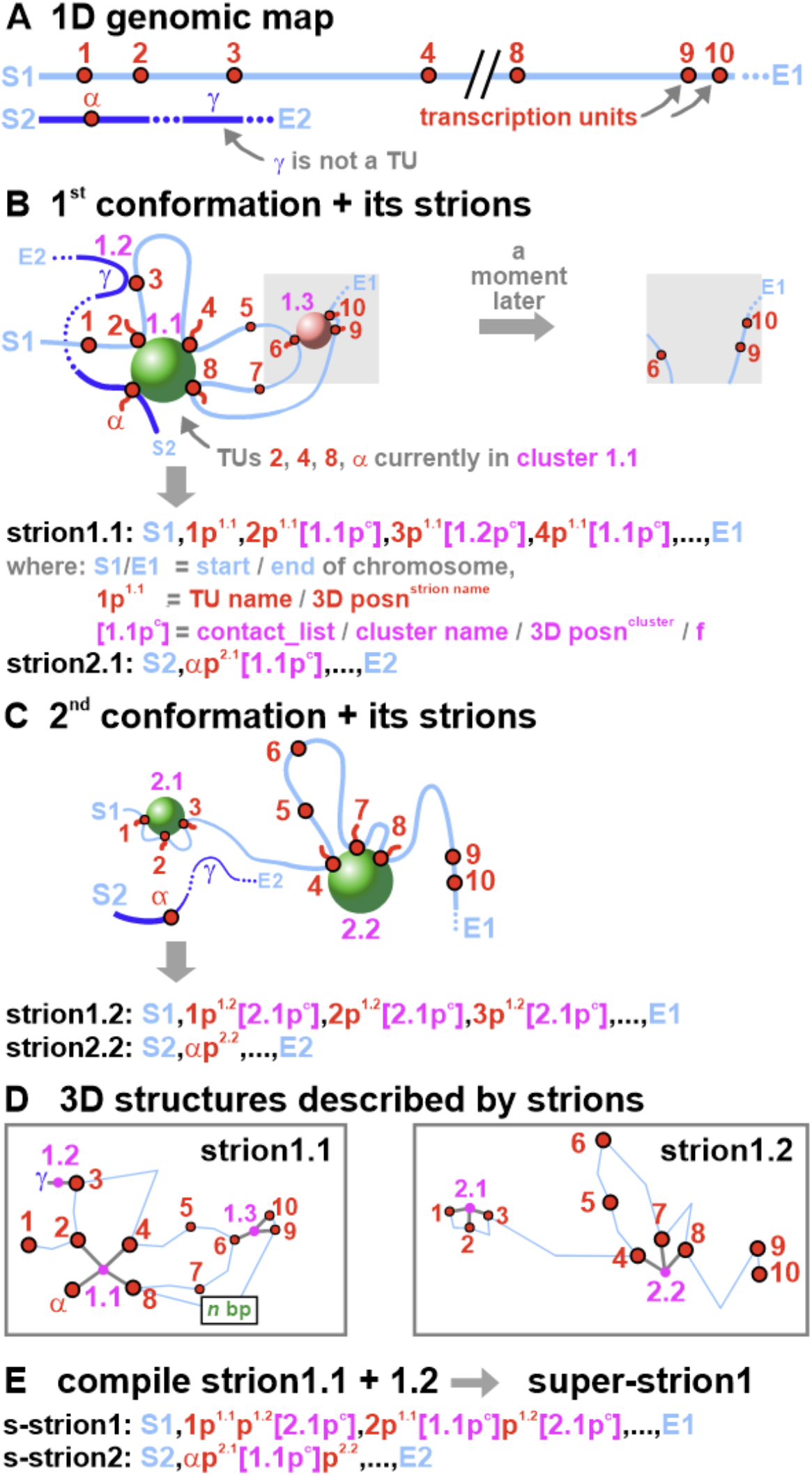
Strions and super-strions. **A**. Map of positions of TUs on example chromosomes 1 and 2. **B**. One conformation these chromosomes might adopt, and their strions. Grey inset shows the same region a moment later, when cluster 1.3 has disappeared. Strions contain many types of information. These includes 1D genomic coordinates of each TU (not shown here), 3D coordinates of each TU (e.g., p^1.1^) and 3D coordinates of cluster centers (p^c^). They also include c_lists and firing frequencies. For example, in cluster 1.1, TUs 2, 4, 8, and α are in the c_list and all fire (f values not shown here). **C**. A 2^nd^ conformation and its strions. TUs 4 and 8 are the only ones seen together in both conformations. **D**. 3D structures determined by strions1.1 and 1.2. Grey lines radiating from cluster points indicate TUs in c_lists. Blue lines indicate the general chromosome path, with segment length between TUs reflecting the number of base pairs, *n* (but not bp position; to simplify images, *n* is shown only between TUs 8 and 9 in strion1.1). Note that γ is not a TU but is included through its association with TU3. **E**. Super-strions (s-strions) for each chromosome are compiled from strions. Decompiling s-strion1 yields the two structures in (D).

### TU clusters capture critical information concerning structure and function

Now consider two of the myriad of conformations that might be adopted by segments of two imaginary chromosomes in two different diploid cells of one cell type (**Fig. 2A**). In the 1^st^ conformation, we start at the beginning of maternal chromosome 1 (S1) and scan down the chromosome. We name the 1^st^ cluster seen as 1.1. This cluster is shown as one green sphere in **Figure 2B**; it contains many components that continually exchange with the soluble pool so that the cluster might remain—very roughly—in the same place for some time before disappearing completely. This cluster anchors four active TUs: 2, 4, 8 (all on chromosome 1) plus *α* (on maternal chromosome 2). The 2^nd^ cluster seen on chromosome 1 (named 1.2) involves a (chance) *trans*-contact between currently inactive TU3 (on chromosome 1) and non-TU *γ* (on chromosome 2). The 3^rd^ cluster (named 1.3) is shown pink because it contains 3 co-transcribed TUs that bind the same (pink) factors. In this case, the three TUs in cluster 1.3 might unbind a moment later; consequently, the cluster disappears, and now-detached TUs join the unstructured fraction (**Fig. 2B**, grey inset).

In the 2^nd^ cell examined, the conformation is likely to be different (**Fig. 2C**). However, some TUs may be transcribed in both conformations. Here, TUs 4 and 8 are co-transcribed in clusters 1.1 and 2.2. They will constitute part of a (green) small-world network if seen together in later cells. Our key assumption is that the four TU clusters (1.1, 1.3, 2.1, and 2.2)—against a background illustrated by cluster 1.2—determine the structure and transcriptional activity of these two chromosomal segments at the particular moment chosen for sampling (reviewed by Negro *et al*., 2024).

### Strions

The central line of code in our database is a ‘strion’—a *stri*ng capturing *str*ucture and funct*ion* (here, transcriptional activity) of each chromosome. Each strion encodes a single conformation of one chromosome by listing each TU, which cluster (if any) to which that TU belongs, and the TUs activity at that moment.

Strion1.1 describes the first conformation of maternal chromosome 1 (**Fig. 2B**), strion1.2 the second (**Fig. 2C**), and so on. Each strion begins with ‘S’ (start) and ends with ‘E’ (end). In between, successive TUs are listed from first to last. Each TU has an unique name (1,2, … 10 …, *α*, …), 1D genomic coordinates (e.g., bases 1, 111-2, 222; not shown here), and a 3D position (marked by p^strion name^). If a TU is in a cluster, this is depicted by ‘[cluster name]’ (purple in **Fig. 2**). Each cluster has an unique bipartite name. The 1^st^ cluster seen in the 1^st^ conformation associated with chromosome 1 is called 1.1, the 2^nd^ is called 1.2; and so on for this and other chromosomes. When a cluster anchors distant parts of one chromosome or even a different chromosome, it keeps its first-used name. For example, cluster 1.1 is first named with respect to TU2 and retains this name when associated with TU4 on chromosome 1 and TUα on chromosome 2. Four types of information are associated with each cluster: name, 3D position (p^c^), c_list and firing frequency (f; values not shown in Figures or text). Strion1.1 begins with TU1 and its 3D position, indicated by ‘S1,1p^1.1^’ (**Fig. 2B**). Next comes TU2 with the same kinds of information plus its associated cluster, summarized by ‘2p^1.1^[1.1p^c^]’. TU3 is in cluster 1.2 (with *γ*) and is described by ‘3p^1.1^[1.2p^c^]’. Note that α and *γ* on chromosome 2 are parts of cluster 1.1 and 1.2 respectively. The first previously-unseen cluster on chromosome 2 will be called 2.1.

Key features of strions include the following. Strions are concise because they discard 3D positional information of disordered regions out in loops, except for that concerning TUs. However, critical positional information is preserved: the 1D and 3D coordinates of each TU on a chromosome, plus the 3D locations of all clusters in each individual conformation. Consequently, strions1.1 and 1.2 define the two chromosome structures illustrated in **Figure 2D**, where blue lines reflect the general (imprecise) paths of chromosomes in space, and line lengths the loop lengths. A strion also contains information on (*trans*) contacts made with other chromosomes through shared clusters. In **Figure 2B**, chromosomes 1 and 2 contact each other through two pathways: TUs 2 and *α* in cluster 1.1, plus TUs 3 and *γ* in cluster 1.2. Two critical functional features are preserved, represented by the purple ‘[‘ and ‘]’ bracketing each cluster in strions. The first is the c_list—a list of all TUs in a cluster. The second allows determination of the firing frequency, f, of each TU. Thus, f of any TU is proportional to the number of times that TU is seen in a cluster across the set of conformations analyzed (Brackley *et al*., 2021; Chiang *et al*., 2022; Negro *et al*., 2024). In its simplest form, f = 1 if a TU is in a cluster (and so ‘fires’), and f = 0 if not (and so is not transcribed). In the latter case, no ‘[‘ or f value are included in a strion. We also define F as the average of all f values for that TU across all cells sampled. F then reflects F’, the transcriptional activity of a TU determined from the GRO-seq reads for that TU relative to the total number of reads of all TUs in the population sampled. Comparison of F with experimentally-derived F’ then provides a sanity check. Values of F’ can be included in another column in our database, and even replace f values of 0 and 1—but they are not required inputs. Note also that F-value equivalents can be calculated using a simple formula with only one variable—the distance in base-pairs between a TU and its nearest promoter (Negro *et al*., 2024). We suggest strions capture essential features of chromosome structure and function, despite containing relatively few 3D coordinates.

A set of strions for one chromosome—obtained from different cells and/or the same cell at different times—can be compiled into one line of code to give a ‘*s*uper-strion’ (s-strion). Thus, strions1.1 + 1.2 give s-strion1, and 2.1 + 2.2 give s-strion2 (**Fig. 2E**). In s-strion1, TU1 has its 3D positions in the two conformations indicated by ‘p^1.1^’ plus ‘p^1.2^’, and its association with cluster 2.1 in the second conformation by ‘[2.1p^c^]’. TU2 is associated with clusters 1.1 and 2.1, and this double association is indicated by ‘[1.1p^c^]’ plus ‘[2.1p^c^]’. Like strions, s-strions contain information on contacts made with other chromosomes. Super-strions also contain critical features of function (through f, F, or F’) and participation in small-world networks (through c_lists). Consequently, one s-strion consists of one line of code that defines the range of structure and function sampled for one chromosome—whether from different cells or one cell at different times. Thus, s-strion1 can be de-compiled to yield the two structures illustrated in **Figure 2D**.

Optional filtering steps can be applied to remove clusters likely to arise from chance contacts. For example, clusters may be retained only if observed in multiple cells or if they contain a minimum number of TUs. These thresholds can be tuned according to dataset size and resolution. We call filterered strions “strionfs” for short. **Figure 3** summarizes steps in deriving strions and s-strions from a set of experiments on one cell type. Note that 46 s-strions define the structures of all chromosomes in a sampled set of diploid cells of one type.

**Figure 3.**
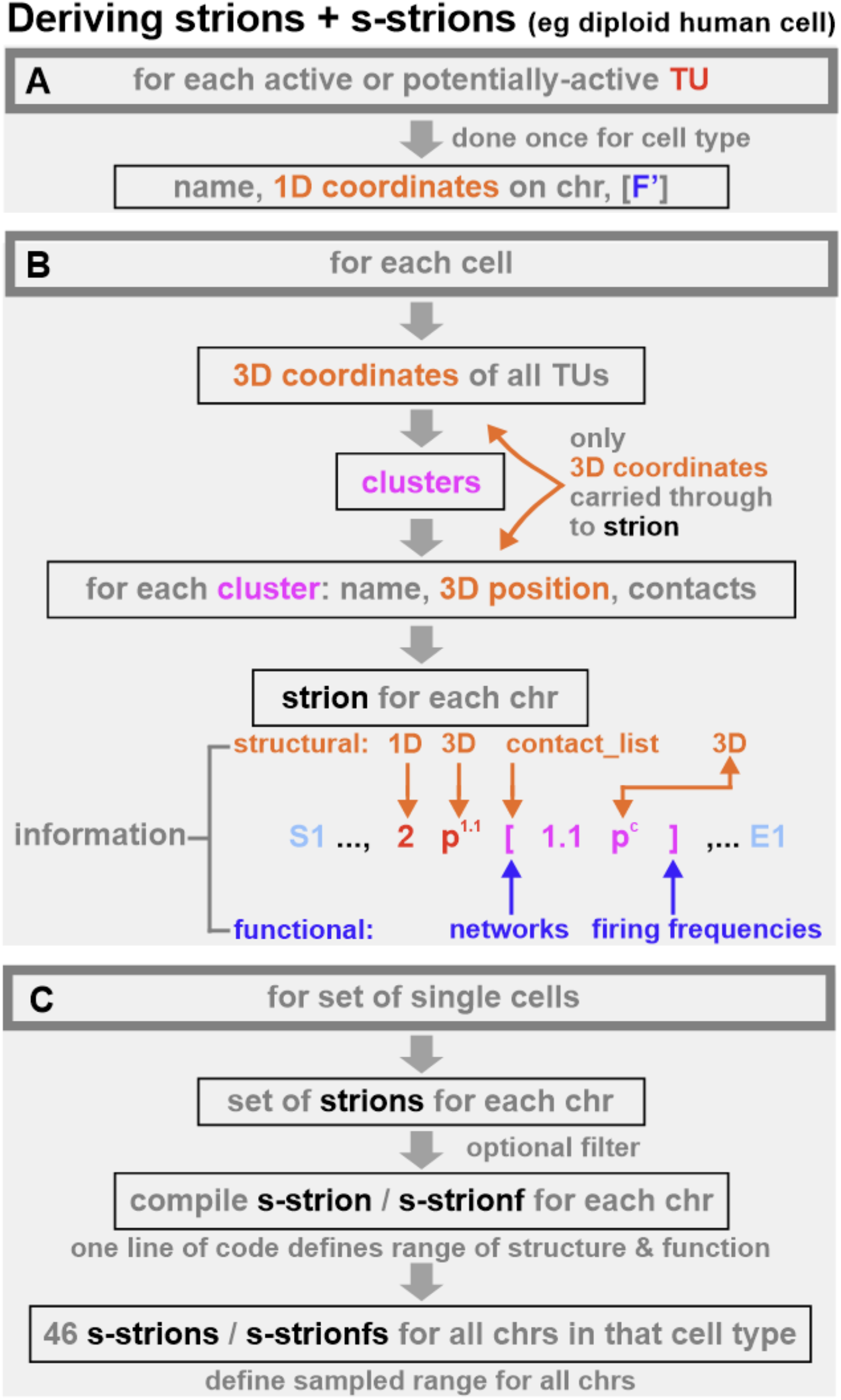
Deriving strions and super-strions. chr: chromosome. **A**. Information required for each active or potentially-active TU. Inclusion of F’ (obtained from GRO-seq data) is optional. **B**. Steps for each cell. The part of strion1.1 involving TU2 is reproduced from Figure 2B and contains the types of information shown (f values are omitted for simplicity but reflect firing frequencies). **C**. Steps for a set of single cells.

The saving in storage requirement associated with strion databases can be demonstrated by the following simple calculation. A typical 50 kDa protein in the PDB takes up 5 Mbytes. Storing a single 100 Mbp chromosome (simulated as 1000, 200, and 1 bp segments) would require 2 Mbytes, 10 Mbytes and 2 Gbytes respectively. Instead, storing a single configuration for a 100 Mbp chromosome as a strion would require 20 Kbytes of memory—a massive compression of information.

### Example strion database

To illustrate the practical implementation of this formalism, we generated a publicly-available database of strions derived from HiP-HoP polymer simulations of human chromosomes in hESC and HUVEC cells (https://git.ecdf.ed.ac.uk/dmarendu/3dgenome_strion_database/). In these examples, each conformation is encoded as simple arrays describing TU cluster membership together with transcriptional state (active/inactive), providing concrete instances of how strions compactly capture structure and function. This database demonstrates how strions can be constructed using data from simulations, and it is easy to imagine how data derived using other methods might be incorporated into it.

The databases also contains simple (toy) strions, which describe chromatin loop networks arising in simpler polymer models for chromatin organisation, such as described in (Bonato et al., 2024; Bonato et al., 2025); more generally, a mathematical discussion of words related to graphs (which chromatin loop networks essentially are) is given in (Kitaev and Lozin, 2015).

### Some uses of s-strions

S-strions provide a compact representation of chromosome structure and transcriptional activity across cells or time points. They can be mined to establish recurrent structural and functional patterns, including for example, loops, boundaries between TADs (topologically associating domains; Lieberman-Aiden *et al*., 2009), and the frequency of *cis* and *trans* contacts made by individual TUs. Note that spurious cluster associations arising from chance contacts can be reduced using filters like the ones we suggest. We now describe some other features of interest.

#### Firing frequencies

An f value of unity is assigned if a TU is in a cluster and zero if not—with F providing the average. Thus, in the two conformations in **Figure 2**, F values for TUs 1, 2, and 5 are respectively (0+1)/2 = 0.5, (1+1)/2 = 1, and (0+0)/2 = 0 (as TU1 is in a cluster only in the 2^nd^ conformation, TU2 is in a cluster in both, and TU5 is never in a cluster). Then, F values are in rank order: TU2 > TU1 > TU5, and we have seen these should reflect ranks of F’-values obtained from GRO-seq.

#### Small-world networks

S-strions reveal small-world networks by showing which TUs co-cluster most frequently. For example, TUs 4 and 8 are in a cluster in both conformations in **Figure 2** (i.e., in cluster 1.1 and 2.2), and this association is carried through into the s-strion. If seen together in more conformations, they are likely part of a small-world (green) network. Moreover, this network can be expanded to include TUs like 2 and *α* (both are in cluster 1.1 with TUs 4 and 8), plus TUs 1 and 3 (both in cluster 2.1 with TU2), if they often co-cluster in later cells examined.

#### Structural and functional motifs

Proteins are often built using structural modules like α-helices and β-sheets, and gene-expression networks are built using functional modules like feed-forward loops and bi-fans (Milo *et al*., 2002). Chromosomes likely have analogous building blocks that our database can help identify.

#### Positive and negative correlations of transcriptional activity

In the conformation in **Figure 2B**, transcription of TUs 9 and 10 are correlated—they are co-active and co-inactive in the left- and right-hand grey zones. Consequently, correlations between any pair or group of TUs can be determined to give co-firing and co-bursting frequencies, for example.

#### Enhancer, silencer, and eQTL interactomes

Enhancers work by being transcribed in close contact with their target genes (Andersson *et al*., 2014; Furlong and Levine, 2018; Schoenfelder and Fraser, 2019; Goel *et al*., 2023; Negro *et al*., 2024). Therefore, TUs that cluster frequently with a gene define that gene’s enhancer interactome (He *et al*., 2014; Andersson and Sandelin, 2020; Negro *et al*., 2024). We suggest that silencers and expression quantitative loci (eQTLs) work much like enhancers—as active TUs that contact their targets (Negro *et al*., 2024). If so, analogous silencer and eQTL interactomes can be derived. We conclude that s-strions support a wide range of structural and functional analyses, even though they contain the 3D coordinates of <4% of the genome (**Fig. 4A**).

**Figure 4.**
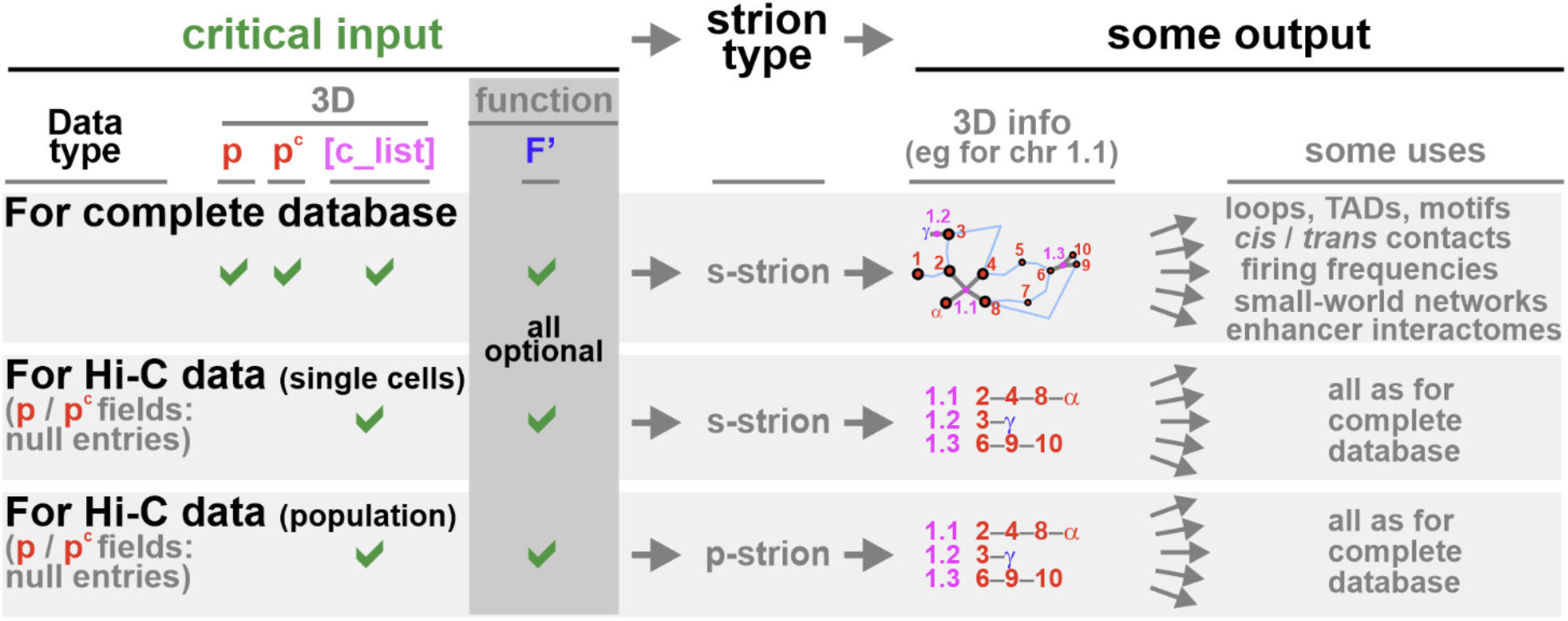
Comparing inputs and outputs for different data sets (info: information). Examples are from Figure 2. With all data sets, F’ is an optional input (as an f value is always automatically assigned if a TU is in a cluster). With the complete database, decompiling an s-strion yields a partial structure and many uses. With current Hi-C data, decompiling an s-strion or p-strion yields the same set of uses, though without 3D coordinates.

### Databases lacking 3D coordinates capture critical structural and functional information

The highest-resolution Hi-C datasets currently available provide extensive contact information across large cell populations, but do not directly include 3D coordinates of chromosomal segments (e.g., Harris *et al*., 2023). Single-cell Hi-C data are sparser and similarly lack explicit spatial coordinates (Zhang R *et al*., 2022; Zhang S *et al*., 2022). This raises a key question: how useful are strion-based representations in the absence of 3D information?

The answer is that they remain highly informative. To understand this answer, consider **Figure 2E** and the part of strion1.1 that lists some properties of TU2 (i.e., ‘2p^1.1^[1.1p^c^c_list]’). This part contains three types of structural information: the 1D genomic coordinates of TU2, 3D coordinates given by p^1.1^ plus p^c^, and the contact list “c_list” (which gives all contacts seen in the cluster, but without 3D coordinates). While p^1.1^ and p^c^ allow us to locate TU2 and its cluster in 3D space, it is the genomic coordinates, c_list, and associated f, F, or F’ values that determine all the useful features described above. These features alone are sufficient to recover structural and functional relationships described above.

For single-cell Hi-C data, strions can therefore be constructed as before, but with null entries for 3D coordinates (**Fig. 4**). In this case, a strion encodes the linear arrangement of TUs together with their clustering relationships. Aggregating these into s-strions retains information on contact frequencies and co-clustering patterns across cells, enabling the same downstream analyses as in the complete database, albeit without explicit spatial reconstruction.

For population-based Hi-C data, a related strategy is adopted (**Fig. 4**). However, we can no longer define strions, as there is no single-cell data. Nevertheless, analogs of s-strions for each chromosome can still be constructed. We call these population-strions (p-strions); they can be thought of as pre-compiled versions derived from a population. As for single-cell Hi-C data, p and p^c^ fields have null entries, optional filtering steps can be included, and the p-strion database retains all uses of the complete one through 1D genomic coordinates, f, F, or F’ values, and c_lists.

## Conclusions

In summary, we have discussed the type of information one would like to see in a database of chromosome structures. We have also proposed symbols and syntax suitable for both the present and the future, when 3D positional information is available for every bp in the genome. In this framework, a *strion*—a single line of code—encodes the path of a chromosome as it passes through 3D space in one cell. Critically, strions encode 3D coordinates of genomic segments or bases separately from inter-segmental contacts and firing frequencies. Thus, strions retain utility even when 3D coordinates are unavailable. Sets of strions for that chromosome are now compiled into an s-strion—again a single line of code—that defines chromosome paths in each cell in the sampled set (**Fig. 2**).

By focusing on TUs and their clustering, strions and s-strions provide a highly compressed representation that discards most positional information from disordered regions while retaining the key features required to describe chromosome organisation and activity. These include contact patterns, loop anchors, chromosome territories, and functional properties such as firing frequencies, enhancer interactomes, and small-world transcriptional networks (**Fig. 4**). Importantly, this representation remains informative even in the absence of explicit 3D coordinates, allowing current datasets from Hi-C, pore-C, GAM, SPRITE, RCMC, and imaging to be incorporated directly.

Strions and s-strions are adaptable in other ways. They are easily integrated with PDB-like databases of atomic structures to add functional information. We focused on TUs and clusters, but other motifs (e.g., transcription start sites, promoters, ATAC-seq peaks, histone/CTCF/cohesin marks) and landmarks like centromeres, nucleoli, and contacts with lamins can be added. It is also straightforward to imagine how strions and s-strions might capture different functions; for example, the frequency and timing of origin firing, as replication also occurs in clusters (Cook, 1999; Marchal *et al*., 2019). In the long term, strions may provide for chromosome conformations a role partially analogous to that played by SMILES strings in cheminformatics (Weininger, 1988): a compact, standardised, machine-readable syntax for storage, comparison, and analysis.

Our proposed databases have inherent drawbacks. It deliberately discards large amounts of data and does not aim to represent any single instantaneous chromosome configuration. Instead, it captures recurring structural and functional features across a dynamic ensemble. As such, its utility will depend on whether the retained information proves sufficient to explain genome organisation and activity at the scales of interest.

We therefore view this work as a starting point for discussion. The central challenge for the future will be to define representations that are both sufficiently compact to handle rapidly-growing datasets and sufficiently flexible to accommodate the unexpected, and hence capture the principles governing chromosome structure and function.

## Acknowledgements

We thank the Wellcome Trust (223097/Z/21/Z) for financial support.

## References

Arrastia, M.V., Jachowicz, J.W., Ollikainen, N., Curtis, M.S., Lai, C., Quinodoz, S.A., Selck, D.A., Ismagilov, R.F., and Guttman, M. (2022). Single-cell measurement of higher-order 3D genome organization with scSPRITE. Nature Biotechnol. 40, 64–73. 10.1038/s41587-021-00998-1

Andersson, R., Gebhard, C., Miguel-Escalada, I., Hoof, I., Bornholdt, J., Boyd, M., Chen, Y., Zhao, X., Schmidl, C., Suzuki, T., et al. (2014). An atlas of active enhancers across human cell types and tissues. Nature 507, 455–461. 10.1038/nature12787

Andersson, R., and Sandelin, A. (2020). Determinants of enhancer and promoter activities of regulatory elements. Nature Rev. Genet. 21, 71–87. 10.1038/s41576-019-0173-8

Badia-i-Mompel, P., Wessels, L., Müller-Dott, S., Trimbour, R., Ramirez Flores, R.O., Argelaguet, R., and Saez-Rodriguez, J. (2023). Gene regulatory network inference in the era of single-cell multi-omics. Nature Rev. Genet. 24, 739–754. 10.1038/s41576-023-00618-5

Beagrie, R.A., Scialdone, A., Schueler, M., Kraemer, D.C.A., Chotalia, M., Xie, S.Q., Barbieri, M., de Santiago, I., Lavitas, L-M., Branco, M.R., et al. (2017). Complex multi-enhancer contacts captured by genome architecture mapping. Nature 543, 519–524. 10.1038/nature21411

Beagrie, R.A., Thieme, C.J., Annunziatella, C., Baugher, C., Zhang, Y., Schueler, M., Kukalev, A., Kempfer, R., Chiariello, A.M., Bianco, S., Li Y., et al. (2023). Multiplex-GAM: genome-wide identification of chromatin contacts yields insights overlooked by Hi-C. Nature Methods 20, 1037–1047. 10.1038/s41592-023-01903-1

Berman, H.M., Westbrook, J., Feng, Z., Gilliland, G., Bhat, T.N., Weissig, H., Shindyalov, I.N., and Bourne, P.E. (2000). The Protein Data Bank. Nucl. Acids Res., 28, 235–242. 10.1093/nar/28.1.235

Bianco, S. Lupiáñez, D.G., Chiariello, A.M., Annunziatella, C., Kraft, K., Schöpflin, R., Wittler, L., Andrey, G., Vingron, M., Pombo, A., Mundlos S., and Nicodemi, M. (2018). Polymer physics predicts the effects of structural variants on chromatin architecture. Nat. Genet. 50, 662–667. 10.1038/s41588-018-0098-8

Bignaud, A., Cockram, C., Borde, C., Groseille, J., Allemand, E., Thierry, A., Marbouty, M., Mozziconacci, J., Espéli, O., and Koszul, R. (2024). Transcription-induced domains form the elementary constraining building blocks of bacterial chromosomes. Nature Structural & Molecular Biology 31, 489–497. 10.1038/s41594-023-01178-2

Bintu, B., Mateo, L.J., Su, J.-H., Sinnott-Armstriong, N.A., Parker, M., Kinrot, S., Yamaya, K., Boettiger, A.N., and Zhuang, X. (2018). Super-resolution chromatin tracing reveals domains and cooperative interactions in single cells. Science 362, eaau1783. https://10.1126/science.aau17

Bonato, A., Chiang, M., Corbett, D., Kitaev, S., Marenduzzo, D., Morozov, A., and Orlandini, E. (2024). Topological spectra and entropy of chromatin loop networks, Phys. Rev. Lett. 132, 248403, 10.1103/PhysRevLett.132.248403

Bonato, A., Carlon, E., Kitaev, S., Marenduzzo, D., and Orlandini, E. (2025). Topological weight and structural diversity of polydisperse chromatin loop networks, 2507.00520, 10.48550/arXiv.2507.00520

Brackley, C. A., Gilbert, N., Michieletto, D., Papantonis, A., Pereira, M.C., Cook, P.R., and Marenduzzo, D. (2021). Complex small-world regulatory networks emerge from the 3D organisation of the human genome. Nat. Commun. 12, 5756. 10.1038/s41467-021-25875-y

Chiang, M., Brackley, C.A., Marenduzzo, D., and Gilbert, N. (2022). Predicting genome organisation and function with mechanistic modelling. Trends Genet. 38, 364–378. 10.1016/j.tig.2021.11.001

Chiang, M., Brackley, C.A., Naughton, C., Nozawa, R.S., Battaglia, C., Marenduzzo, D., and Gilbert, N. (2024). Genome-wide chromosome architecture prediction reveals biophysical principles underlying gene structure. Cell Genomics 4, 100698. 10.1016/j.xgen.2024.100698

Contessoto, V.G., Cheng, R.R., Hajitaheri, A., Dodero-Rojas, E., Mello, M.F., Lieberman-Aiden, E., Wolynes, P.G., Di Pierro, M., and Onuchic, J.N. (2021). The Nucleome Data Bank: web-based resources to simulate and analyze the three-dimensional genome. Nucleic Acids Res. 49(D1), D172–D182. 10.1093/nar/gkaa818

Cook, P.R. (1999). The organization of replication and transcription. Science 284, 1790–1795. 10.1126/science.284.5421.1790

Cook, P.R., and Brazell, I.A. (1975). Supercoils in human DNA. J. Cell Sci 19, 261–279. 10.1242/jcs.19.2.261

Cook, P.R., and Marenduzzo, D. (2018). Transcription-driven genome organization: a model for chromosome structure and the regulation of gene expression tested through simulations. Nucl. Acids Res. 46, 9895–9906. 10.1093/nar/gky763

Dekker, J., et al. (2026). An integrated view of the structure and function of the human 4D nucleome. Nature 649, 759–776. 10.1038/s41586-025-09890-3

Deshpande, A.S., Ulahannan, N., Pendleton, M., Dai, X., Ly, L., Behr, J.M., Schwenk, S., Liao, W., Augello, M.A., Tyer, C., et al. (2022). Identifying synergistic high-order 3D chromatin conformations from genome-scale nanopore concatemer sequencing. Nature Biotechnol. 40, 1488–1499. 10.1038/s41587-022-01289-z

Di Pierro, M., Potoyan, D.A., Wolynes, P.G., and Onuchic, J.N. (2018). Anomalous diffusion, spatial coherence, and viscoelasticity from the energy landscape of human chromosomes. Proc. Natl. Acad. Sci. U.S.A. 115, 7753–7758. 10.1073/pnas.1806297115

Dotson, G.A., Chen, C., Lindsly, S., Cicalo, A., Dilworth, S., Ryan, C., Jeyarajan, S., Meixner, W., Stansbury, C., Pickard, J., et al. (2022). Deciphering multi-way interactions in the human genome. Nature Commun. 13, 5498. 10.1038/s41467-022-32980-z

Fullwood, M.J., Liu, M.H., Pan, Y.F., Liu, J., Xu, H., Mohamed, Y.B., Orlov, Y.L., Velkov, S., Ho, A., Mei, P.H., et al. (2009). An oestrogen-receptor--bound human chromatin interactome. Nature, 462, 58–64. 10.1038/nature08497

Furlong, E.E.M., and Levine, M. (2018). Developmental enhancers and chromosome topology. Science 361, 1341–1345. 10.1126/science.aau0320

Giorgetti, L., Galupa, R., Nora, E.P., Piolot, T., Lam, F., Dekker, J., Tiana, G., and Heard, E. (2014). Predictive polymer modeling reveals coupled fluctuations in chromosome conformation and transcription. Cell. 157, 950–963. 10.1016/j.cell.2014.03.025

Goel, V.Y., Huseyin, M.K., and Hansen, A.S. (2023). Region Capture Micro-C reveals coalescence of enhancers and promoters into nested microcompartments. Nature Genet. 55, 1048–1056. 10.1038/s41588-023-01391-1

Harris, H.L., Gu, H., Olshansky, M., Wang, A., Farabella, I., Eliaz, Y., Kalluchi, A., Krishna, A., Jacobs, M., et al. (2023). Chromatin alternates between A and B compartments at kilobase scale for subgenic organization. Nature Commun. 14, 3303. 10.1038/s41467-023-38429-1

Hatton, I.A., Galbraith, E.D., Merleau, N.S.C., Miettinen, T.P., Smith, B.M., and Shander, J.A. (2023). The human cell count and size distribution. Proc, Natl. Acad. Sci. USA 120, e2303077120. 10.1073/pnas.2303077120

He, B., Chen, C., Teng, Li., and Tan, K. (2014). Global view of enhancer–promoter interactome in human cells. Proc. Natl. Acad. Sci. U.S.A. 111, E2191–E2199. 10.1073/pnas.1320308111

HuBMAP Consortium (2019). The human body at cellular resolution: the NIH Human Biomolecular Atlas Program. Nature 574, 187–192. 10.1038/s41586-019-1629-x

Jackson, D.A., Dickinson, P., and Cook, P.R. (1990). The size of chromatin loops in HeLa cells. EMBO J. 9, 567–571. 10.1002/j.1460-2075.1990.tb08144.x

Jordán-Pla, A., Pérez-Martínez, M.E., and Pérez-Ortín, J.E. (2019). Measuring RNA polymerase activity genome-wide with high-resolution run-on-based methods. Methods 159-160, 177–182. 10.1016/j.ymeth.2019.01.017

Kempfer, R., and Pombo, A. (2020). Methods for mapping 3D chromosome architecture. Nature Rev. Genet. 21, 207–226. 10.1038/s41576-019-0195-2

Kitaev, S., and Lozin, V. (2015). Words and graphs. Springer International Publishing. https://link.springer.com/book/10.1007/978-3-319-25859-1

Kwak, H., Fuda, N.J., Core, L.J., and Lis, J.T. (2013). Precise maps of RNA polymerase reveal how promoters direct initiation and pausing. Science 339, 950–953. 10.1126/science.1229386

Larsson, A.J.M., Johnsson, P., Hagemann-Jensen, M., Hartmanis, L., Faridani, O.R., et al. (2019). Genomic encoding of transcriptional burst kinetics. Nature 565, 251–254. 10.1038/s41586-018-0836-1

Lerner, J., Gomez-Garcia, P.A., McCarthy, R.L., Liu, Z., Lakadamyali, M., and Zaret, K.S. (2020). Two-parameter mobility assessments discriminate diverse regulatory factor behaviors in chromatin. Mol. Cell 79, 677-688.e6. 10.1016/j.molcel.2020.05.036

Li, G.L., Ruan, X.A., Auerbach, R.K., Sandhu, K.S., Zheng, M.Z., Wang, P., Poh, H.M., Goh, Y., Lim, J., Zhang, J.Y., et al. (2012). Extensive promoter-centered chromatin interactions provide a topological basis for transcription regulation. Cell 148, 84–98. 10.1016/j.cell.2011.12.014

Li, W., Lu, J., Lu, P., Gao, Y., Bai, Y., Chen, K., Su, X., Li, M., Liu, J., Chen, Y., et al. (2023). scNanoHi-C: a single-cell long-read concatemer sequencing method to reveal high-order chromatin structures within individual cells. Nature Methods 20, 1493–1505. 10.1038/s41592-023-01978-w

Lieberman-Aiden, E., Van Berkum, N.L., Williams, L., Imakaev, M., Ragoczy, T., Telling, A., Amit, I., Lajoie, B.R., Sabo, P.J., Dorschner, M.O. et al. (2009). Comprehensive mapping of long-range interactions reveals folding principles of the human genome. Science 326, 289–293. 10.1126/science.1181369

Marchal, C., Sima, J., and Gilbert, D.M. (2019). Control of DNA replication timing in the 3D genome. Nature Revs Mol. Cell Biol. 20, 721–737. 10.1038/s41580-019-0162-y

Misteli, T. (2020). The self-organizing genome: principles of genome architecture and function. Cell 183, P28–45. 10.1016/j.cell.2020.09.014

Nagano, T., Várnai, C., Schoenfelder, S., Javierre, B.-M., Wingett, S.W., and Fraser, P. (2015). Comparison of Hi-C results using in-solution versus in-nucleus ligation. Genome Biol. 16, 175. 10.1186/s13059-015-0753-7

Negro, G., Semeraro, M., Cook, P.R., and Marenduzzo, D. (2024). A unified-field theory of genome organization and gene regulation. iScience 24, 111218. 10.1016/j.isci.2024.111218

Nickerson, J. (2001). Experimental observations of a nuclear matrix. J. Cell Sci. 114, 463–474. 10.1242/jcs.114.3.463

Nir, G., Farabella, I., Estrada, C.P., Ebeling, C.G., Beliveau, B.J., Sasaki, H.M., Lee, S.H., Nguyen, S.C., McCole, R.B., Chattoraj, S., et al. (2018). Walking along chromosomes with super-resolution imaging, contact maps, and integrative modeling. PLoS Genet. 14, e1007872. 10.1371/journal.pgen.1007872

Olivares-Chauvet, P., Mukamel, Z., Lifshitz, A., Schwartzman, O., Elkayam, N.O., Lubling, Y., Deikus, G., Sebra, R.P., and Tanay, A. (2016). Capturing pairwise and multi-way chromosomal conformations using chromosomal walks. Nature 540, 296–300. 10.1038/nature20158

Oluwadare, O., Highsmith, M., and Cheng, J. (2019). An overview of methods for reconstructing 3-D chromosome and genome structures from Hi-C data. Biol. Proced. Online 21, 7. 10.1186/s12575-019-0094-0

Oluwadare, O., Highsmith, M., Turner, D., Lieberman Aiden, E., and Cheng. J. (2020). GSDB: a database of 3D chromosome and genome structures reconstructed from Hi-C data. BMC Mol. Cell Biol. 21, 60. 10.1186/s12860-020-00304-y

O’Sullivan, J.M., Hendy, M.D., Pichugina, T., Wake, G.C.G., and Langowski, J. (2013). The statistical-mechanics of chromosome conformation capture. Nucleus 4, 390–398. 10.4161/nucl.26513

Papantonis, A., Kohro, T., Baboo, S., Larkin, J., Deng, B., Short, P., Tsutsumi, T., Taylor, S., Kanki, Y., Kobayashi, et al. (2012). TNFα signals through specialized factories where responsive coding and micro-RNA genes are transcribed. EMBO J. 31, 4404–4414. 10.1038/emboj.2012.288

Pombo, A., Jackson, D.A., Hollinshead, M., Wang, Z., Roeder, R.G., and Cook, P.R. (1999).Regional specialization in human nuclei: visualization of discrete sites of transcription by RNA polymerase III. EMBO J. 18, 2241–2253. 10.1093/emboj/18.8.2241

Pownall, M.E., Miao, L., Vejnar, C.E., M’Saad, O., Sherrard, A., Frederick, M.A., Benitez, M.D.J., Boswell, C.W., Zaret, K.S., et al. (2023). Chromatin expansion microscopy reveals nanoscale organization of transcription and chromatin. Science 381, 92–100. 10.1126/science.ade5308

Quinodoz, S.A., Ollikainen, N., Tabak, B., Palla, A., Schmidt, J.M., Detmar, E., Lai, M.M., Shishkin, A.A., Bhat, P., Takei, Y., et al. (2018). Higher-order inter-chromosomal hubs shape 3D genome organization in the nucleus. Cell 174, 744–757. 10.1016/j.cell.2018.05.024

Rippe, K. and Papantonis, A. (2025) RNA polymerase II transcription compartments — from factories to condensates. Nature Revs. Genet. 26, 775–788. 10.1038/s41576-025-00859-6

Rowley, M.J., Poulet, A., Nichols, M.H., Bixler, B.J., Sanborn, A.L., Brouhard, E.A., Hermetz, K., Linsenbaum, H., Csankovszki, G., Aiden, E.L., et al. (2020). Analysis of Hi-C data using SIP effectively identifies loops in organisms from C. elegans to mammals. Genome Res. 30, 447–458. 10.1101/gr.257832.119

Schoenfelder, S., and Fraser, P. (2019). Long-range enhancer-promoter contacts in gene expression control. Nature Rev. Genet. 20, 437–455. 10.1038/s41576-019-0128-0

Schoenfelder, S., Sexton, T., Chakalova, L., Cope, N.F., Horton, A., Andrews, S., Kurukuti, S., Mitchell, J.A., Umlauf, D., Dimitrova, D.S. et al. (2010). Preferential associations between co-regulated genes reveal a transcriptional interactome in erythroid cells. Nature Genet., 42, 53–61. 10.1038/ng.496

Semeraro, M., Negro, G., Forte, G., Suma, A., Gonnella, G., Cook, P.R., and Marenduzzo, D. (2025). Cluster size determines morphology of transcription factories in human cells. eLife 14, RP103955. 10.7554/eLife.103955.2

Shah, S., Takei, Y., Zhou, W., Lubeck, E., Yun, J., Eng, C.-H.L., Koulena, N., Cronin, C., Karp, C., Liaw, E.J., et al., (2018). Dynamics and spatial genomics of the nascent transcriptome by intron seqFISH. Cell 174, 363–376. 10.1016/j.cell.2018.05.035

Tiana, G., and Giorgetti, L. (2018). Integrating experiment, theory and simulation to determine the structure and dynamics of mammalian chromosomes. Curr. Opin. Struct. Biol. 49, 11–17. 10.1016/j.sbi.2017.10.016

van Noort, V., Snel, B., and Huynen, M.A. (2004). The yeast coexpression network has a small-world, scale-free architecture and can be explained by a simple model. EMBO Rep. 5, 280–284. 10.1038/sj.embor.7400090

Weininger, D. (1988). SMILES, a chemical language and information system. 1. Introduction to methodology and encoding rules. J. Chem. Inf. Comput. Sci. 28, 31–36. https://pubs.acs.org/doi/10.1021/ci00057a005

Winick-Ng, W., Kukalev, A., Harabula, I., Zea-Redondo, L., Szabó, D., Meijer, M., Serebreni, L., Zhang, Y., Bianco, S., Chiariello, A.M., et al. (2021). Cell-type specialization is encoded by specific chromatin topologies. Nature 599, 684–691. 10.1038/s41586-021-04081-2

Yildirim, A., Boninsegna, L., Zhan, Y., and Alber, F. (2022). Uncovering the principles of genome folding by 3D chromatin modeling. Cold Spring Harbor Perspect. Biol. a039693v1. 10.1101/cshperspect.a039693

Zhang, R., Zhou, T., and Ma, J. (2022). Multiscale and integrative single-cell Hi-C analysis with Higashi. Nature Biotechnol. 40, 254–261. 10.1038/s41587-021-01034-y

Zhang, S., Plummer, D., Lu, L., Cui, J., Xu, W., Wang, M., Liu, X., Prabhakar, N., Shrinet, J., Srinivasan, D., et al. (2022). DeepLoop robustly maps chromatin interactions from sparse allele-resolved or single-cell Hi-C data at kilobase resolution. Nature Genet. 54, 1013–1025. 10.1038/s41588-022-01116-w

Zhang, S., Übelmesser, N., Barbieri, M., and Papantonis, A. (2023). Enhancer–promoter contact formation requires RNAPII and antagonizes loop extrusion. Nature Genet. 55, 832–840. 10.1038/s41588-023-01364-4

